# STAM Interaction with Hrs Controls JAK/STAT Activation by Interferon-α at the Early Endosome

**DOI:** 10.1101/509968

**Authors:** Natacha Zanin, Cédric M. Blouin, Christine Viaris de Lesegno, Daniela Chmiest, Ludger Johannes, Christophe Lamaze

## Abstract

Activation of the Janus kinase (JAK)/signal transducer and activator of transcription (STAT) pathway by type I interferons (IFN) requires clathrin-dependent endocytosis of the IFN-α/β receptor (IFNAR). The molecular machinery that brings about the selective activation of IFN-α/β-induced JAK/STAT signaling on endosomes remains unknown. Here we show that the constitutive association of STAM with IFNAR1 and the TYK2 Janus kinase at the plasma membrane prevents the activation of TYK2 by type I IFNs. IFN-α stimulated endocytosis leads to the interaction of IFNAR1 with Hrs on early endosomes, which then relieves TYK2 inhibition by STAM and thereby allows for TYK2 and IFNAR signaling. In contrast, IFN-β stimulation results in sorting of IFNAR to a distinct endosomal subdomain where the receptor is activated independently from Hrs. Our results identify the molecular machinery that controls the spatiotemporal activation of TYK2 and establish the central role of endosomal sorting in the differential regulation of JAK/STAT signaling by IFN-α and IFN-β.

**Summary:** The spatiotemporal activation of JAK/STAT signaling by IFN-α is controlled by STAM association with Hrs at the early endosome.

## INTRODUCTION

The four members of the Janus kinase (JAK) family of tyrosine kinases (JAK1, JAK2, JAK3 and TYK2) play a major role in cell signaling as signal downstream transducers of more than fifty cytokine and growth factor receptors, and are associated with a large spectrum of human diseases (*1*). How this limited set of tyrosine kinases can control the activation, regulation and pleiotropic functions of such a vast family of signaling receptors remains obscure. This question is particularly salient for type I interferons (IFNs) where sixteen human IFNs transduce their signal through binding to a common cell surface heterodimeric receptor, IFNAR, made of one dimer of IFNAR1 and IFNAR2 subunit, respectively associated with TYK2 and JAK1 (*2*). Numerous investigations have tried to understand how IFN**-**α and IFN**-**β can mediate different signaling outputs and physiological activities through the activation of the same JAK/signal transducer and activator of transcription (STAT) pathway. Seminal studies on epidermal growth factor receptor (EGF-R) endocytosis and signaling have contributed to define the concept of the signaling endosome, where the spatiotemporal propagation and amplification of signaling can be actively controlled (*3-6*). The current challenge is to identify the molecular machineries that control signaling at the endosome. Our finding that clathrin-dependent endocytosis of the IFNAR complex was required for the activation of JAK/STAT signaling by IFN**-**α suggested a possible role for endosomal sorting in JAK/STAT activation (*7*). Here, we identify the molecular machinery that controls the spatiotemporal activation of JAK/STAT signaling induced by IFN-α at the early endosome.

## RESULTS

### IFNAR1 interacts with Hrs at the early endosome

We monitored by immunofluorescence the intracellular fate of the IFNAR complex after 10 min of endocytosis in cells stimulated with IFN-α or IFN-β stimulation. We found that the majority of the endocytosed IFNAR1 subunits were present in early endosomes, as shown by colocalization with the *bona fide* endosomal marker EEA1 (Fig. 1A). These endosomes were also positive for hepatocyte growth factor-regulated tyrosine kinase substrate (Hrs), a member of the endosomal sorting complex required for transport (ESCRT)-0 that binds to ubiquitylated cargo proteins and sorts them to the degradative lysosomal pathway (*8-10*). Whereas IFNAR1 and Hrs immunostainings were perfectly superimposed upon cell stimulation with IFN-α, we found that for IFN-β, IFNAR1 staining was juxtaposed to Hrs, suggesting that IFNAR was localized in a different endosomal subdomain.

**Fig. 1.**
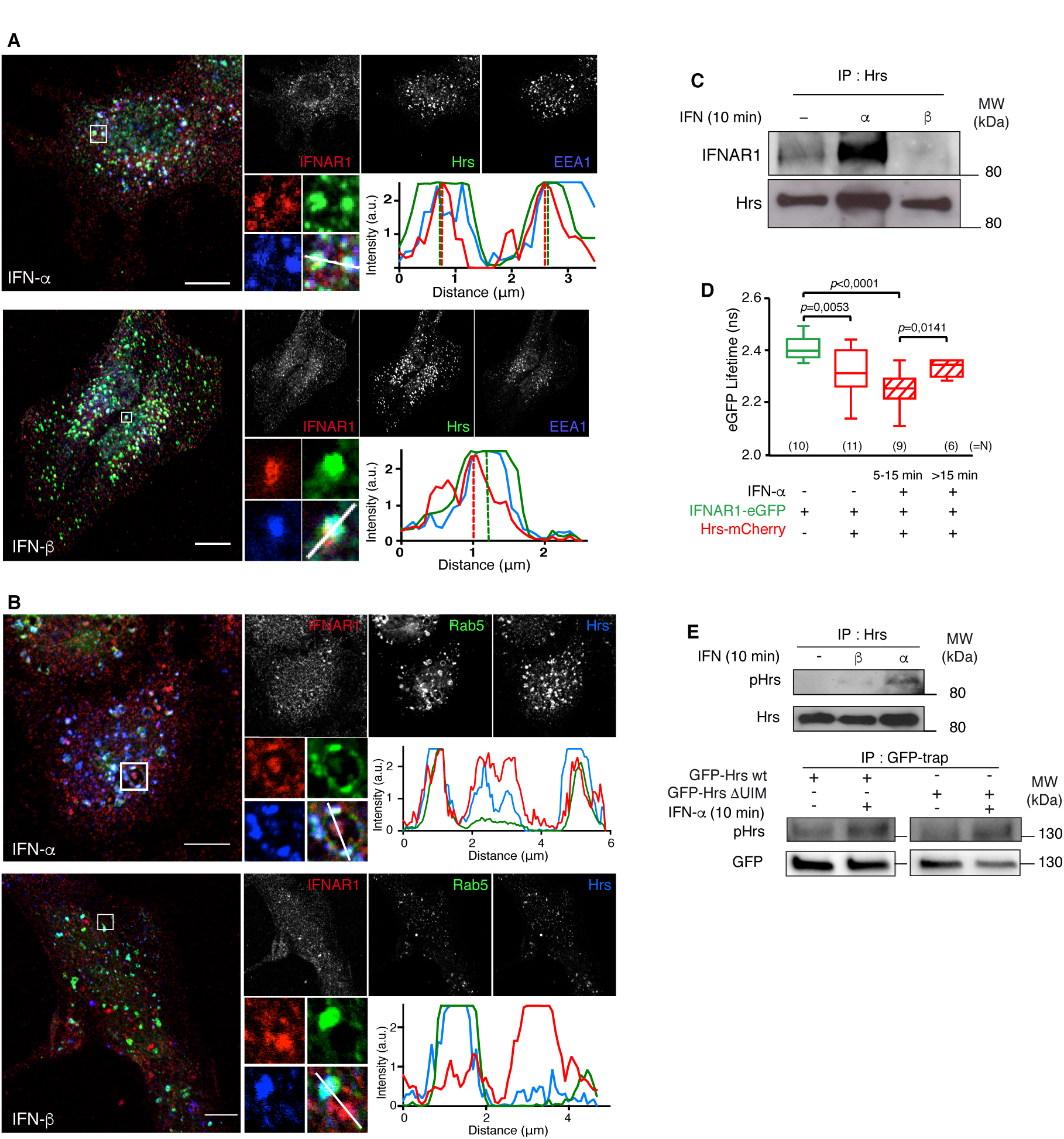
IFNAR1 interaction with Hrs. (**A**) Immunofluorescent labeling of Hrs, EEA1 and endocytosed IFNAR1 subunits after 10 min IFN-α (upper panels) or IFN-β (lower panels) stimulation observed by confocal microscopy. Representative plots of colocalization profiles on endosomes. Scale bar 10 µm. (**B**) Immunofluorescent labeling of Hrs and endocytosed IFNAR1 subunits in eGFP-Rab5 Q79L-transfected RPE1 cells after 10 min IFN-α (upper panel) or IFN-β (lower panel) stimulation observed by confocal microscopy. Representative plots of colocalization profiles on endosomes (lower panels). Scale bar 10 µm. (**C**) RPE1 cells were stimulated or not (-) with IFN-α or IFN-β for 10 min as indicated. Cell lysates were incubated with anti-Hrs antibody to immunoprecipitate (IP) endogenous Hrs and reveal co-immunoprecipitated endogenous IFNAR1. (**D**) Kinetics of IFNAR1 interaction with Hrs measured by FLIM-FRET during IFN-α stimulation in RPE1 cells expressing Hrs-mCherry and/or IFNAR1-eGFP as indicated. P-values were calculated with Mann-Whitney Test. (**E**) Tyrosine phosphorylation of Hrs (pHrs) was revealed in Hrs immunoprecipitates from RPE1 cells stimulated or not (-) with IFN-α or IFN-β for 10 min as indicated (upper panel). Tyrosine phosphorylation of Hrs was revealed in GFP-trap immunoprecipitates from RPE1 cells transfected with wild type (wt) eGFP-Hrs or with eGFP-Hrs ΔUIM deletion mutant (unable to bind to ubiquitin) and stimulated or not (-) with IFN-α or IFN-β for 10 min as indicated (lower panel). **A**-**E** are representative of 3 independent experiments.

This important aspect was further analyzed on enlarged endosomes that form upon expression of the GTPase-deficient Rab5 mutant Q79L (*37*). On such enlarged endosomes, we could clearly observe that the distribution of internalized IFNAR1 subunits differed with the IFN subtype (Fig. 1B). After 10 min of stimulation with IFN-α, the endocytosed IFNAR1 subunits fully colocalized with Hrs and Rab5. In contrast, after IFN-β stimulation, the IFNAR1 subunits were localized in distinct endosomal subdomains devoid of Hrs and Rab5 (Fig. 1B).

By immunoprecipitation experiments, it was found that IFNAR1 and Hrs interact (Fig. 1C). This interaction was minimal in unstimulated cells and dramatically increased after 10 min of cell stimulation by IFN-α. In contrast, we could not detect IFNAR1 subunits in Hrs immunoprecipitates from cells stimulated by IFN-β (Fig. 1C).

We next combined fluorescence lifetime imaging (FLIM) and Förster resonance energy transfer (FRET) microscopy to monitor in live cells the molecular dynamics of the interaction of IFNAR1 with Hrs at high spatial and temporal resolution. At steady state, IFNAR1-eGFP fluorescence lifetime decreased when Hrs-mCherry was co-expressed, confirming the physical interaction between IFNAR1 and Hrs (Fig. 1D). This effect was strongest after 5 to 15 min IFN-α stimulation, reflecting an increased interaction with Hrs at the early endosome (*8*). Later than 15 min of IFN-α stimulation, IFNAR1-eGFP fluorescence lifetime increased again significantly, which is consistent with the sorting of IFNAR1 subunits to late endosomes where Hrs is no longer present (*11*).

Hrs was initially characterized as a tyrosine phosphorylated substrate of several growth factors, especially hepatocyte growth factor (*8-13*). We found that a 10 min stimulation by IFN-α significantly increased Hrs tyrosine phosphorylation over basal levels found in unstimulated cells. In contrast, no Hrs tyrosine phosphorylation could be detected when cells were stimulated by IFN-β (Fig. 1E). It is classically described that Hrs interacts with ubiquitinylated cargo proteins through its ubiquitin interacting motif (UIM) (*14, 15*). A mutated form of Hrs, deleted from the UIM, presented a similar increase of tyrosine phosphorylation after stimulation by IFN-α (Fig. 1E), suggesting that IFNAR1 ubiquitination was not required for Hrs interaction and phosphorylation, as already reported for other cytokine receptors (*16*). This result is also consistent with our finding that only IFN-α and not IFN-β promotes the interaction of IFNAR1 with Hrs, whereas both IFN-α and IFN-β can induce the ubiquitylation of IFNAR1 subunits (*17, 18*). Altogether these results demonstrate that IFN-α stimulation promotes the ubiquitin-independent phosphorylation of Hrs, and its interaction with IFNAR at the early endosome. In contrast, upon IFN-β stimulation, Hrs is not phosphorylated and does not interact with IFNAR.

### JAK/STAT activation by IFN-α requires Hrs

IFN-mediated signaling relies predominantly on the canonical JAK/STAT signaling pathway (*19*). IFN-α/β binding to IFNAR2 results in IFNAR1 and IFNAR2 subunit dimerization, followed by auto/trans-phosphorylation and activation of the TYK2 and JAK1 tyrosine kinases, which are, respectively, pre-associated with IFNAR1 and IFNAR2 (*20-22*). JAK-mediated IFNAR1 tyrosine phosphorylation allows the recruitment and activation of cytoplasmic STAT1 and STAT2, which in association with IRF9, are translocated to the nucleus as the ISGF3 complex that binds to IFN-stimulated response elements in DNA to initiate gene transcription (*23*). Our finding that clathrin-dependent endocytosis of IFNAR controls the activation of JAK/STAT signaling by type I IFNs (*7*) suggested that the activation of IFNAR-associated JAK kinases could occur at the endosome.

We therefore explored the role of IFNAR binding to Hrs in JAK/STAT signaling. As expected, IFN-α stimulation led to full activation (i.e. tyrosine phosphorylation) of JAK1 and TYK2 (Fig. 2A). In Hrs depleted cells, virtually no activation of JAK1 and TYK2 was detected after 10 min of IFN-α stimulation (Fig. 2A). Hrs depletion did not change the overall levels of intracellular and cell surface IFNAR1 (Fig. S1A). Consistent with the lack of JAK1 and TYK2 activation, the tyrosine phosphorylation of IFNAR1, a requisite for STAT1 recruitment, could not be detected in the absence of Hrs (Fig. 2A). As a result, IFN-α-stimulated tyrosine phosphorylation of STAT1 (pSTAT1) was strongly inhibited in Hrs depleted cells (Fig. 2B). Impaired STAT1 activation in Hrs-depleted cells translated into an absence of pSTAT1 translocation to the nucleus (Fig. S1B). In agreement with the lack of IFNAR interaction with Hrs under IFN-β stimulation conditions, Hrs depletion did not alter JAK/STAT activation by IFN-β (Fig. 2B and S1B). Supporting our finding that Hrs activation by IFN-α occurs independently from its UIM domain (Fig. 1E), mouse embryonic fibroblasts (MEF) expressing human IFNAR2 and a human IFNAR1 mutant deleted for its ubiquitination sites, displayed nevertheless sustained JAK/STAT signaling in response to IFN-α stimulation (Fig. 2C).

**Fig. 2.**
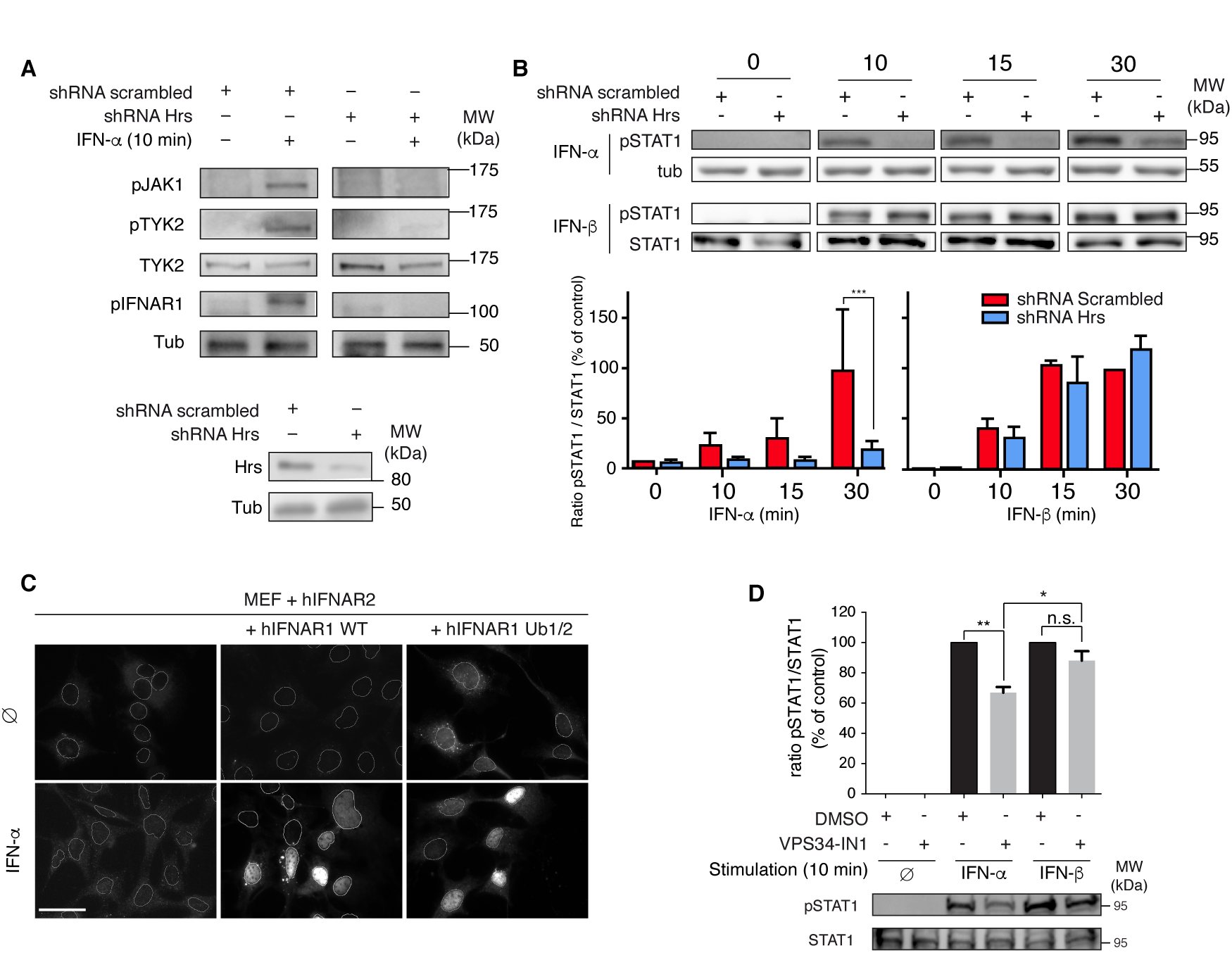
IFNAR1 interaction with Hrs is required for IFN-α-induced JAK/STAT signaling. (**A**) Immunoblots for tyrosine phosphorylation levels of JAK1 (pJAK1), TYK2 (pTYK2) and IFNAR1 (pIFNAR1) in control (shRNA scrambled) and Hrs depleted (shRNA Hrs) RPE1 cells stimulated or not for 10 min with IFN-α as indicated. Lower panel shows the extent of shRNA-mediated Hrs depletion. (**B**) Immunoblots for tyrosine phosphorylation levels of STAT1 (pSTAT1) in control (shRNA scrambled) and Hrs depleted (shRNA Hrs) RPE1 cells stimulated or not with IFN-α or IFN-β for the indicated times. Data are expressed as the pSTAT1/STAT1 ratio for each condition, the shRNA scrambled/IFN-β 30 min condition being set at 100 %. Data represent the mean ratio ± SEM of 3 independent experiments. P-values were calculated with two-tailed unpaired T-test. *** P<0.001. (**C**) Nuclear translocation of pSTAT1 in mouse embryonic fibroblasts (MEF) transfected with human IFNAR2 and IFNAR1 wt or the Ub1/2 mutant deleted from the 3 ubiquitination sites. Scale bar 50 µm. (**D**) Immunoblots (lower panel) and quantification (upper panel) of STAT1 tyrosine phosphorylation (pSTAT1) in HeLa cells treated or not with the Vps34 inhibitor (VPS34-IN1) and stimulated for 10 min with IFN-α or IFN-β. The ratio pSTAT1/STAT1 was calculated for each condition, the shRNA scrambled/IFN-β 10 min condition being set at 100 %. Data represent the mean ratio ± SEM of 3 independent experiments. P-values were determined using one-way ANOVA with Dunnett’s multiple comparison test. \**P*<0.05; \*\**P*<0.01; ns, not significant.

Depleting PI3-P levels through pharmacological inhibition of the PI3 kinase Vps34 also led to a specific decrease of STAT1 tyrosine phosphorylation by IFN-α (Fig 2D), in agreement with the recruitment of Hrs to early endosomes through PI3-P binding (*8, 33*). PI3-P depletion did not affect STAT1 activation by IFN-β confirming that endocytosed IFNAR were localized to distinct endosomal subdomains after IFN-β stimulation (Fig. 2D).

### STAM-dependent signaling block at the plasma membrane

We next sought to identify the molecular mechanisms by which Hrs controls IFNAR-dependent JAK/STAT activation at the early endosome. Signal transducing adaptor molecule (STAM) binds tightly to Hrs to form the ESCRT-0 complex at the early endosome (*15, 24, 25*). In humans, the STAM family consists of STAM1, STAM2A and STAM2B, which share conserved domains that are important for their function (*26*). Interestingly, the four members of the JAK family were shown to interact with the ITAM domain present in STAM1 and STAM2A in co-immunoprecipitation experiments (*27, 28*). We studied therefore the possible role of STAM1 and STAM2A in the control of JAK activation by IFNs. We focused our studies on human STAM1 and Hbp, the murine orthologue of human STAM2A (*26*). In agreement with published data (*25, 29*), we found STAM2A mainly localized in early endosomal structures positive for EEA1 (Fig. 3A). At steady state, a fraction of TYK2 was also associated with these EEA1 and STAM2A-positive endosomes. Accordingly, TYK2 significantly co-immunoprecipitated with STAM2A, irrespective of IFN-α and IFN-β stimulation (Fig. S1C). FRET by FLIM measurements on EEA1 positive endosomes confirmed the constitutive interaction of TYK2 with STAM2A independently from IFN-α and IFN-β stimulation (Fig. 3A).

**Fig. 3.**
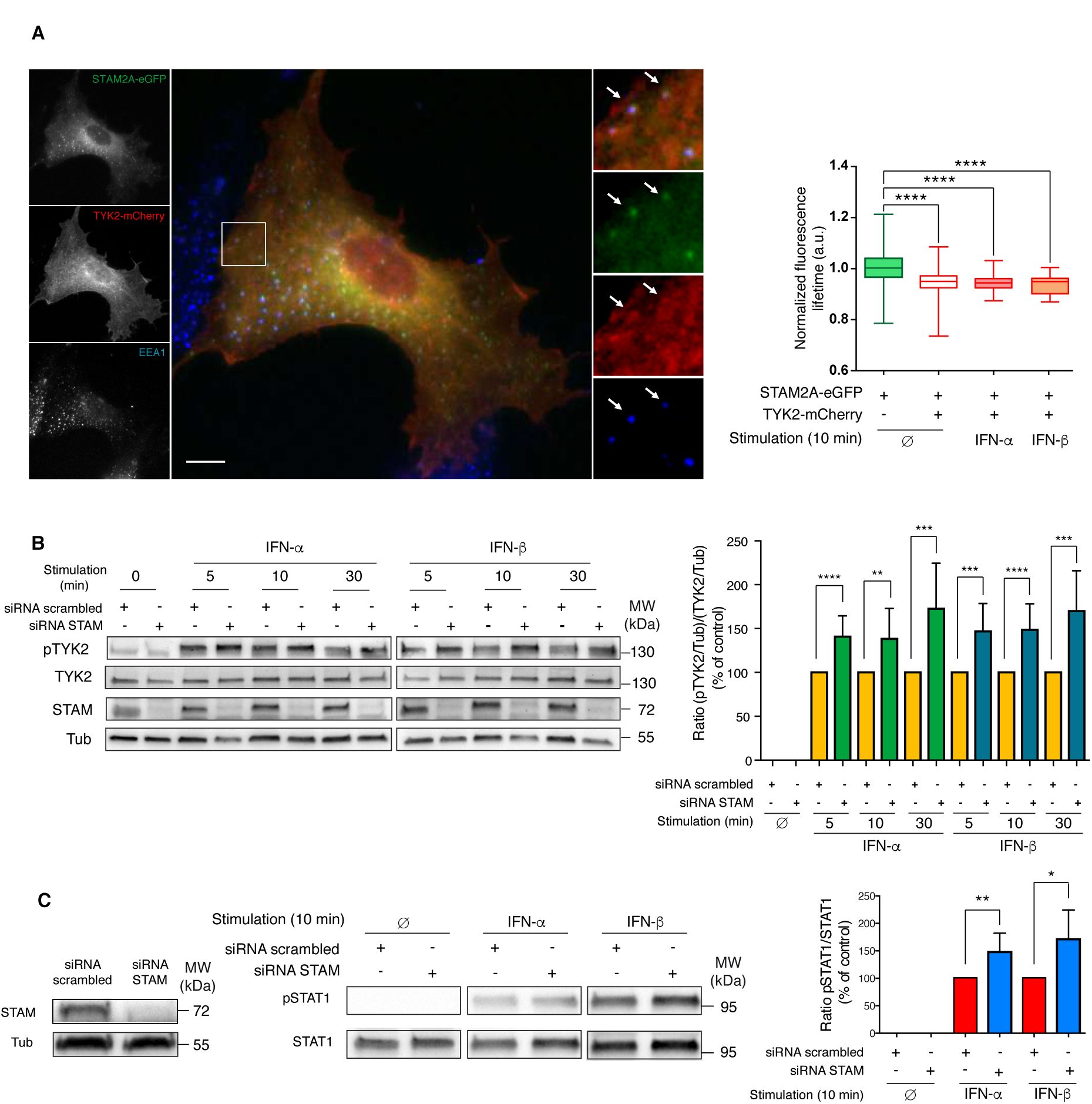
The constitutive association of STAM with TYK2 prevents TYK2 activation. (**A**) Left, widefield microscopy showing STAM2A and TYK2 colocalization at steady state in early endosomes labeled for EEA1 in RPE1 cells transiently transfected with STAM2A-eGFP and TYK2-mCherry. Right, FLIM-FRET measurements in early endosomes reveal a constitutive interaction between STAM2A-eGFP (donor) and TYK2-mCherry (acceptor). Data shown are mean values ± SD of STAM2A-eGFP fluorescence lifetime from 2 independent experiments (normalized). A one-way ANOVA with Tukey’s multiple comparison test was used to compare endosomes stained with STAM2A-eGFP alone (set as control condition) vs. endosomes co-stained with STAM2A-eGFP and TYK2-mCherry. (**B)** Left, kinetics of TYK2 tyrosine phosphorylation (pTYK2) upon IFN-α or IFN-β stimulation in RPE1 cells depleted for STAM. Right, pTYK2 and TYK2 levels were normalized to the loading control level (tubulin), and the ratio (pTYK2/Tub)/(TYK2/Tub) was calculated for each condition and expressed as a percent of control. (**C**) Left panel shows the extent of siRNA-mediated STAM depletion. Middle, STAT1 tyrosine phosphorylation (pSTAT1) levels upon IFN-α or IFN-β stimulation for 10 min in RPE1 cells depleted or not (Ø) for STAM. Right, pSTAT1 level was normalized to STAT1 level and the ratio pSTAT1/STAT1 was calculated for each condition, the siRNA scrambled/IFN-β 10 min condition being set at 100 %. Data in **B** and **C** are representative of 3 to 5 independent experiments. Associated quantifications show mean values ± SEM. Statistical significance was determined using a two-tailed t test. \**P*<0.05; \*\**P*<0.01; \*\*\**P*<0.001; \*\*\*\**P*<0.0001.

We next explored the role of STAM2A in TYK2-mediated signaling. IFN-α or IFN-β-induced tyrosine phosphorylation of TYK2 and STAT1 were increased in STAM1 and STAM2-depleted cells, both at early and later times (Fig. 3B,C). STAM depletion did not change the intracellular and cell surface amounts of IFNAR1 nor the kinetics of IFNAR1 uptake (Fig. S2). These results imply that STAM acts as a negative regulator of JAK/STAT signaling by constitutively inhibiting the TYK2 kinase activity independently from IFN-α or IFN-β stimulation. This is in contrast to Hrs, which is a positive regulator of TYK2 kinase activity and JAK/STAT signaling under IFN-α but not IFN-β, stimulation conditions (Fig. 2A,B).

These opposing effects are difficult to reconcile with the established role of Hrs/STAM as an obligate hetero-dimeric or tetrameric ESCRT-0 complex operating at the early endosome (*15, 30, 31*). We therefore tested the possibility that STAM and Hrs may act independently from each other. Figure 4A shows that endogenous IFNAR1 can be detected in eGFP-STAM2A pull down lysates from cells stimulated or not with IFN-α or IFN-β. The constitutive interaction of IFNAR1 with STAM2A in the absence of IFN stimulation suggested that STAM may bind to IFNAR1 at the plasma membrane. Indeed, cells transiently expressing mCherry-STAM2A and IFNAR1-eGFP showed a significant level of colocalization at the plasma membrane, in addition to the one on early endosomes (Fig. 4B). Similar results were found at steady state in cells expressing endogenous levels of STAM and IFNAR1 (Fig. S3A).

**Fig. 4.**
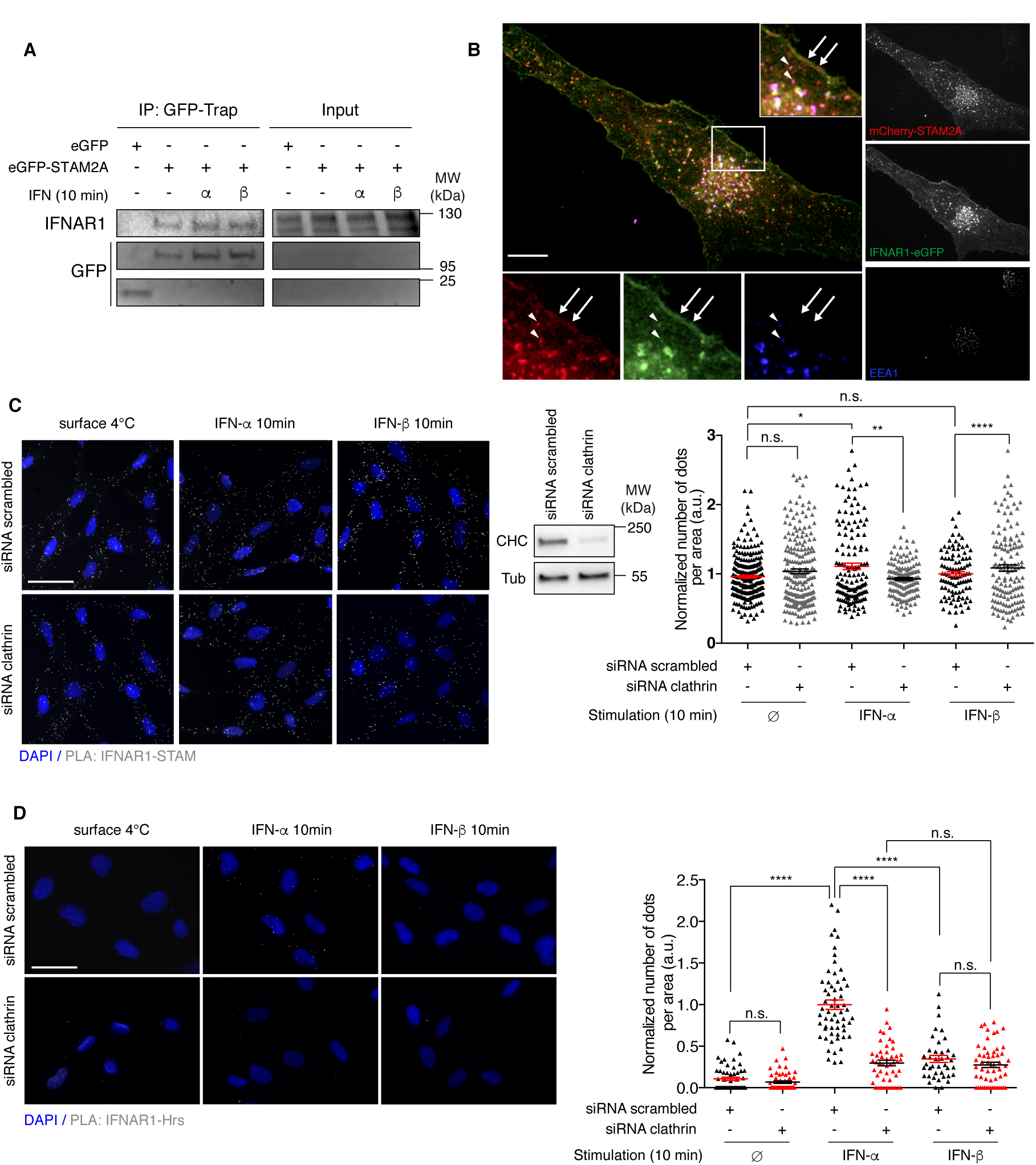
IFNAR1 constitutively interacts with STAM at the plasma membrane and at the early endosome. (**A**) Endogenous IFNAR1 was revealed in GFP-trap immunoprecipitates from RPE1 cells transiently transfected with eGFP-STAM2A and stimulated or not (-) with IFN-α or IFN-β for 10 min as indicated. Representative of 3 independent experiments. (**B**) Widefield microscopy showing co-localization between STAM2A and IFNAR1 at the plasma membrane and early endosomes at steady state in RPE1 cells transiently transfected with mCherry-STAM2A and IFNAR1-eGFP. Scale bar 10 µm. (**C**) Left, *in situ* proximity ligation assay (PLA) experiments monitoring the interaction of surface (4°C) or internalized IFNAR1 with STAM or (**D**) with Hrs after 10 min of IFN stimulation in RPE1 cells (upper panels) or in clathrin-depleted RPE1 cells for 72 h (lower panels). Cells were analyzed by widefield microscopy. Control condition was performed in the absence of primary antibody against STAM or Hrs. Scale bars 50 µm. Nuclei were stained with DAPI and the interaction between the two endogenous proteins of interest was visualized by a dot. Each spot on the graphs in **C** and **D** represents a cell and corresponds to the number of dots in this cell divided by the area of the cell and expressed as arbitrary units (a.u.). Data in **C** and **D** are representative of 3 independent experiments that were normalized for the mean value of the surface condition set at 1 a.u. Corresponding quantifications show mean values ± SEM. Statistical significance was determined using one-way ANOVA with Dunnett’s multiple comparison test. \**P*<0.05; \*\**P*<0.01; \*\*\**P*<0.001; \*\*\*\**P*<0.0001.

We further analyzed the interaction between endogenous STAM and IFNAR1 using the proximity ligation assay (PLA), which allows to directly visualize protein-protein interactions *in situ* (*32*). PLA experiments showed that endogenous STAM and IFNAR1 were found in close proximity at the plasma membrane (within ≈ 40 nm) at steady state, and that this interaction was conserved during IFNAR1 endocytosis under IFN-α or IFN-β stimulation conditions (Fig. 4C). Inhibition of IFNAR1 endocytosis through clathrin depletion confirmed the constitutive interaction of STAM with the pool of IFNAR1 present at the plasma membrane. In agreement with the abovementioned findings (Fig. 1A-D), PLA experiments confirmed the tight interaction of endocytosed IFNAR1 with Hrs after 10 min of IFN-α but not IFN-β stimulation (Fig. 4D). In contrast to STAM however, no constitutive interaction between IFNAR1 and Hrs could be detected at the plasma membrane under resting or IFN-α stimulation conditions, even when IFNAR1 endocytosis was inhibited in clathrin-depleted cells (Fig. 4D). Finally, PLA and co-immunoprecipitation experiments revealed that the depletion of the TYK2 kinase did not prevent IFNAR1 interaction with STAM (Fig. S3B,C). Altogether, these data indicate that at steady state, STAM interacts constitutively with IFNAR1 at the plasma membrane, independently of Hrs and TYK2. Upon IFN-α stimulation and IFNAR1 endocytosis, STAM interaction with IFNAR1 continues at the early endosome where IFNAR1 associates with Hrs.

### IFNAR1 interaction with Hrs relieves STAM inhibition of TYK2

STAM interacts with IFNAR1 at the plasma membrane to control TYK2 activation in an opposite manner to Hrs, whose interaction with IFNAR1 is restricted to the early endosome. This raised the possibility that JAK/STAT signaling was controlled according to the subcellular location of IFNAR. We tested this hypothesis by blocking IFNAR1 endocytosis to the early endosome so as to restrict the analysis of STAM function to the pool of IFNAR present at the plasma membrane. As shown above (Fig. 3C), depletion of STAM led to a significant increase of IFN-α or IFN-β-stimulated STAT1 tyrosine phosphorylation (Fig. 5A). In cells where IFNAR uptake was blocked by clathrin depletion, the activation of STAT1 by either IFN-α or IFN-β was strongly decreased, as reported before (*7*). However, in cells depleted for both clathrin and STAM, we observed a level of STAT1 activation similar or even higher than in control cells (Fig. 5A). Thus, in the absence of STAM, IFNAR endocytosis is dispensable for efficient JAK/STAT signaling. These data suggested that IFNAR endocytosis was required to relieve the inhibitory effect of STAM on TYK2 activity.

**Fig. 5.**
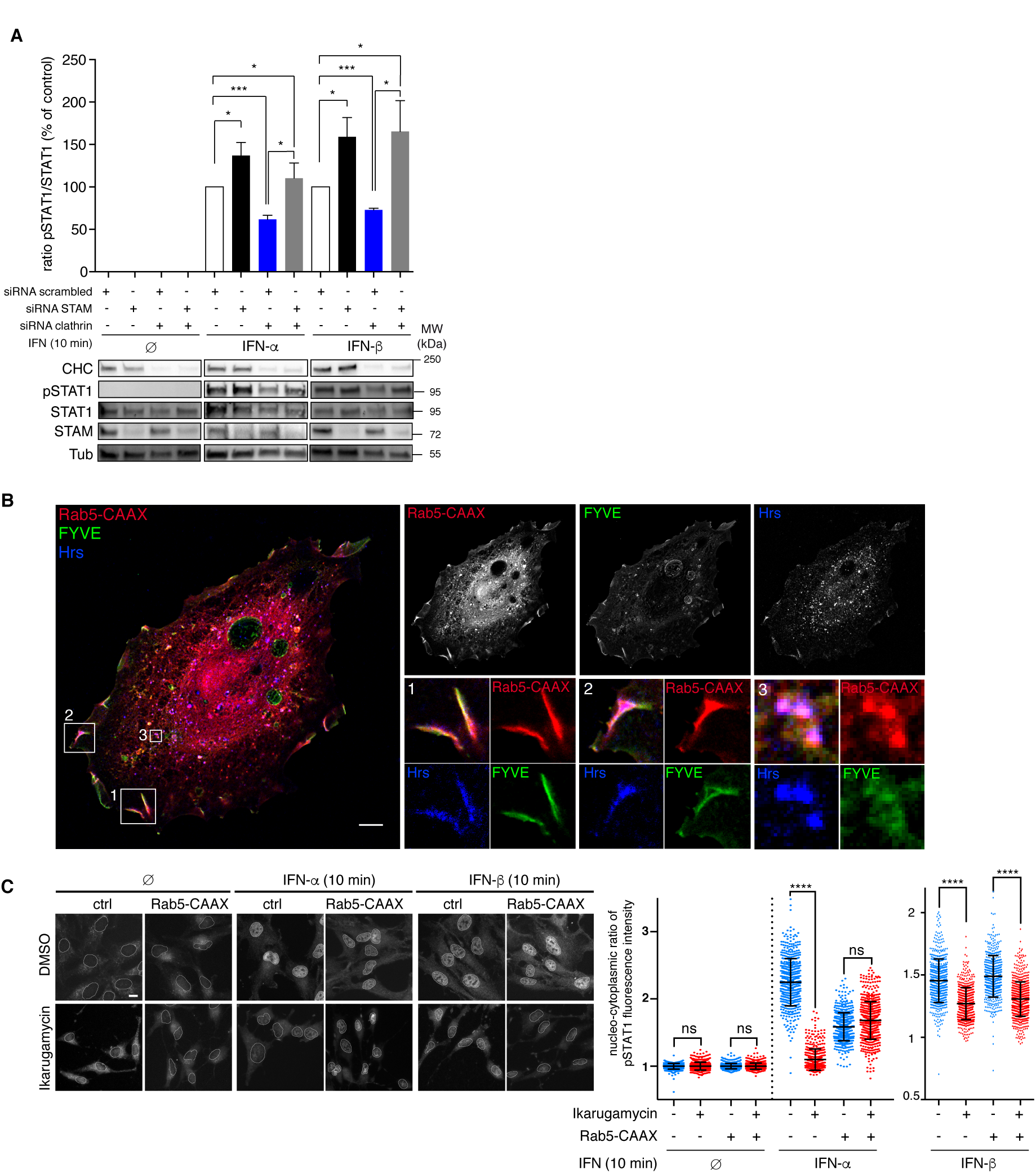
IFNAR1 interaction with Hrs relieves STAM inhibition of TYK2. (**A**) Immunoblots and quantification of STAT1 tyrosine phosphorylation after stimulation of RPE1 cells for 10 min with or without IFN-α or IFN-β in control conditions (white bar), after STAM depletion (black bar), after clathrin depletion (blue bar), and after depletion of both STAM and clathrin (grey bar). pSTAT1 level was normalized to STAT1 level and the ratio pSTAT1/STAT1 was calculated for each condition. Data are representative of 3 to 5 independent experiments and were normalized to scrambled siRNA under IFN-α or IFN-β stimulated conditions. Corresponding quantifications show mean values ± SEM. Statistical significance was determined using one-way ANOVA with Dunnett’s multiple comparison test. \**P*<0.05; \*\**P*<0.01; \*\*\**P*<0.001. (**B**) Immunofluorescent labeling of endogenous Hrs at the plasma membrane (zoom windows 1 and 2) and in endosomal structures (zoom window 3) in RPE1 cells co-transfected with constitutively active RFP-Rab5-CAAX (Q79L) (to target PI3-P production at the plasma membrane) and eGFP-FYVE (to visualize PI3-P domains) observed by confocal microscopy. Scale bar 10 µm. (**C**) pSTAT1 nuclear translocation (left panel) and quantification (right panel) in HA-empty vector (ctrl) or constitutively active HA-Rab5-CAAX (Q79L) transfected RPE1 cells pretreated or not for 20 min at 37°C with 4 μM ikaguramycin (to inhibit clathrin-dependent endocytosis) prior to IFN-α or IFN-β stimulation. The nucleus/cytosol fluorescence ratio for pSTAT1 is quantified from the fluorescence imaging. P-values were calculated with one-way ANOVA with Kruskal-Wallis multiple comparison test. \*\*\*\**P*<0.0001; ns, not significant. Results are representative of 3 independent experiments. Scale bar, 10 µm.

We hypothesized that this would be achieved by the binding of STAM to Hrs at the early endosome. We sought to test this hypothesis by forcing the ectopic localization of Hrs at the plasma membrane. Hrs is recruited to early endosomes through the binding of its FYVE motif to phosphatidylinositol 3-phosphate (PI3-P) (*8, 33*). PI3-P is specifically generated at the early endosome via the phosphorylation of phosphatidylinositol by PI3-kinases, which are recruited and activated at the early endosome by the GTPase Rab5 (*34*). To target constitutively active Rab5 to the plasma membrane, we ectopically expressed the Rab5 Q79L mutant, containing the C-terminal CAAX box of K-Ras (*35*). This resulted in PI3-P production at the plasma membrane, as shown by colocalization between GFP-FYVE (a PI3-P probe) and Rab5-CAAX (Fig. 5B). Under this condition, Hrs also localized at the plasma membrane, in addition to its classical endosomal localization.

To block IFNAR at the plasma membrane, we used ikarugamycin (Ika), a recently characterized natural product inhibitor of clathrin-mediated endocytosis (*36*). In ikarugamycin-treated cells, we indeed detected a strong accumulation of the clathrin pathway cargo transferrin at the cell surface, confirming that the endocytosis block was efficient. A similar effect was observed for IFNAR1 (Fig. S4). As expected (*7*), inhibition of IFNAR endocytosis by ikarugamycin led to a drastic inhibition of STAT1 nuclear translocation in cells stimulated for 10 min by either IFN-α or IFN-β (Fig. 5C). Importantly however, in cells inhibited for IFNAR endocytosis, STAT1 nuclear translocation induced by IFN-α could be rescued if Hrs was ectopically recruited to the plasma membrane. In contrast, localizing Hrs at the plasma membrane did not rescue the nuclear translocation of STAT1 upon stimulation of cells by IFN-β. This finding is consistent with the lack of Hrs involvement in IFNAR sorting and JAK/STAT signaling under IFN-β stimulation conditions (Fig. 1B,C,E and 2B). Hrs was thus able to relieve the inhibition that STAM constitutively exerts on TYK2 activity at the plasma membrane.

## DISCUSSION

It has been more than twenty years since the first report describing the control of epidermal growth factor receptor signaling by endocytosis (*3*). If many receptors, including IFNAR (*7*), were found to follow a similar regulation, the underlying mechanisms remain poorly understood. Here we provide evidence that STAM locks JAK/STAT signaling in a silent mode at the plasma membrane by binding constitutively to IFNAR1 and TYK2. Upon IFN-α stimulation, IFNAR is endocytosed into Hrs and PI3-P positive domains of the early endosome. Binding of Hrs to STAM relieves STAM inhibition on TYK2 and thereby allows the efficient activation of TYK2 by IFN-α at the early endosome (Fig. 6).

**Fig. 6.**
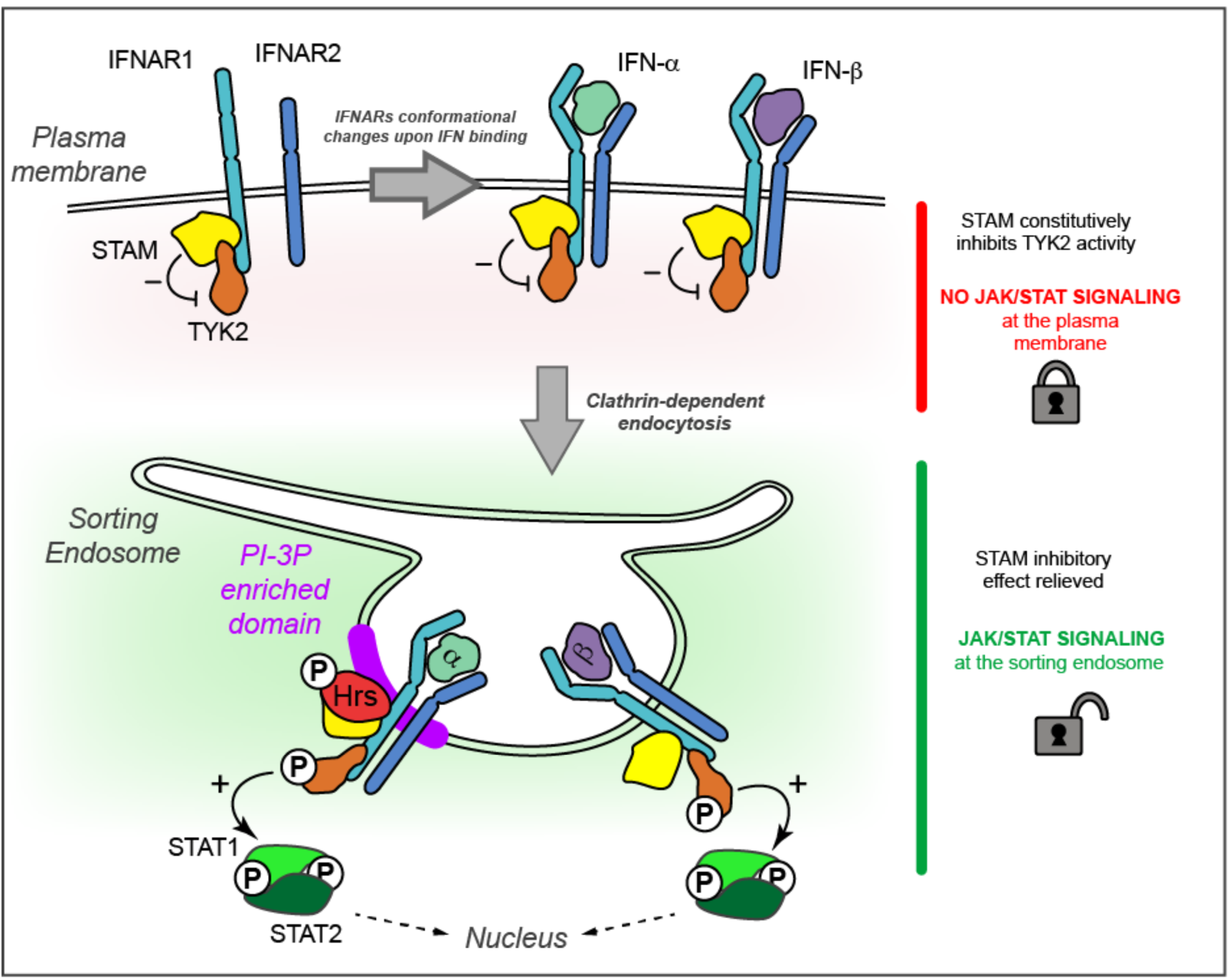
STAM and Hrs control the spatiotemporal activation of JAK/STAT signaling by IFN-α. At steady state, STAM is constitutively associated with IFNAR1 and TYK2 at the plasma membrane where it locks JAK/STAT signaling in a silent mode by inhibiting TYK2 activity. Upon IFN-α stimulation, activated IFNAR is endocytosed by clathrin-dependent endocytosis to the early endosome where it is actively sorted by Hrs into phosphatidylinositol-3-phosphate (PI-3P) enriched domains. By binding to STAM, Hrs relieves the inhibition of TYK2 activity and allows JAK/STAT signaling. In contrast, upon IFN-β stimulation, endocytosed IFNAR1 is delivered to distinct endosomal subdomains devoid of Hrs and PI-3P. Our results establish the central role of Hrs/STAM interaction in the selectivity of JAK/STAT activation by type I interferons at the early endosome.

Our study unveils two unusual facets of STAM and Hrs function. STAM is classically described as a constitutive binding partner of Hrs to assemble ESCRT-0 at the early endosome (*31*). Our results show that STAM can also operate independently from Hrs at the plasma membrane, and that Hrs can be a positive regulator of signaling, in addition to its classical inhibitory action through receptor sorting for lysosomal degradation (*40*).

How specific subtypes of type I IFN can transduce differential responses by binding to the same cell surface receptor has been intensively investigated and remains controversial (*2, 38*). Different affinities toward IFNAR subunits at the plasma membrane were shown to control differential IFN activities by conditioning IFNAR conformational changes, internalization rates, and STAT activation (*21*). In this context, we found that IFN-α2-YNS, an IFN-α variant presenting the same binding affinity to IFNAR as IFN-β (*39*), behaved nevertheless like IFN-α since STAT1 activation by IFN-α2-YNS was still controlled by Hrs (Fig. S1B). Finally, upon IFN-β stimulation, endocytosed IFNAR is targeted to distinct endosomal subdomains devoid of Hrs and PI3-P. These findings establish that in addition to the structural changes occurring at the plasma membrane, IFNAR endosomal sorting is central to the differential activation of JAK/STAT signaling by IFN-α and IFN-β.

Recently, we reported that the retromer complex controls the duration of JAK/STAT signaling by IFN-α and IFN-β through differential sorting of IFNAR1 and IFNAR2 subunits at the early endosome (*41*). We also found that the activation of JAK/STAT signaling by IFN-γ occurred at the plasma membrane through lateral diffusion of the IFN-γR in lipid nanodomains (*42*). Together with the current study, these findings emphasize the critical role of membrane dynamics and trafficking in the differential control of IFN subtype selectivity within the JAK/STAT signaling pathway.

In conclusion, we have uncovered the molecular machinery that allows the endosomal control of signal transduction by IFN-α. The Hrs and STAM interplay that was identified here represents a key switch for TYK2 activity between the plasma membrane and the early endosome. Our study underscores the central role of endosomes as active hubs for the selective control of signaling. While this aspect has long been overlooked for JAK/STAT signaling, future studies should address this regulation in the many pathological contexts where this pathway is altered.

## ACKNOWLEDGEMENTS

The core facilities and the recombinant antibodies platform of Institut Curie as well as the scientific and technical assistance from staff in the PICT-IBiSA / Nikon Imaging Centre at Institut Curie-CNRS, member of the French National Research Infrastructure France-BioImaging (ANR10-INSB-04) are acknowledged. In particular, we would like to thank François Waharte and Lucie Sengmanivong for help with FLIM-FRET and fluorescence microscopy experiments. The help of the following people for providing materials or expertise is acknowledged: Graça Raposo and Philippe Benaroch (Institut Curie, Paris), Pierre Eid (INSERM, Paris). This work was supported by institutional grants from the Curie Institute, INSERM, CNRS, and by specific grants from Agence Nationale de la Recherche (ANR NANOSTAT-15-CE11-0025-01 to CL) and Marie Curie Actions — Networks for Initial Training (FP7-PEOPLE-2010-ITN to CL and DC). NZ was supported by a PhD fellowship from Ministère de l’Enseignement Supérieur et de la Recherche, and Ligue Nationale contre le Cancer, DC by a PhD fellowship from Fondation pour la Recherche Médicale, and CMB by a post-doctoral fellowship from Ligue Nationale contre le Cancer. The Johannes and Lamaze teams are members of Labex CelTisPhyBio N° ANR-10-LBX-0038 part of the IDEX IdexPSL N°ANR-10-IDEX-0001-02 PSL.

## References and Notes

1. J. J. O'Shea et al., The JAK-STAT pathway: impact on human disease and therapeutic intervention. Annual review of medicine 66, 311–328 (2015).

2. J. Piehler, C. Thomas, K. Christopher Garcia, G. Schreiber, Structural and dynamic determinants of type I interferon receptor assembly and their functional interpretation. Immunol Rev 250, 317–334 (2012).

3. A. V. Vieira, C. Lamaze, S. L. Schmid, Control of EGF receptor signaling by clathrin-mediated endocytosis. Science 274, 2086–2089 (1996).

4. P. P. Di Fiore, M. von Zastrow, Endocytosis, signaling, and beyond. Cold Spring Harbor perspectives in biology 6, (2014).

5. R. Irannejad, N. G. Tsvetanova, B. T. Lobingier, M. von Zastrow, Effects of endocytosis on receptor-mediated signaling. Curr Opin Cell Biol 35, 137–143 (2015).

6. R. Villasenor, Y. Kalaidzidis, M. Zerial, Signal processing by the endosomal system. Curr Opin Cell Biol 39, 53–60 (2016).

7. M. Marchetti et al., Stat-mediated Signaling Induced by Type I and Type II Interferons (IFNs) Is Differentially Controlled through Lipid Microdomain Association and Clathrin-dependent Endocytosis of IFN Receptors. Mol Biol Cell. 17, 2896–2909. (2006).

8. S. Urbe, I. G. Mills, H. Stenmark, N. Kitamura, M. J. Clague, Endosomal localization and receptor dynamics determine tyrosine phosphorylation of hepatocyte growth factor-regulated tyrosine kinase substrate. Mol Cell Biol 20, 7685–7692 (2000).

9. S. Urbe et al., The UIM domain of Hrs couples receptor sorting to vesicle formation. J Cell Sci 116, 4169–4179 (2003).

10. P. E. Row, M. J. Clague, S. Urbe, Growth factors induce differential phosphorylation profiles of the Hrs-STAM complex: a common node in signalling networks with signal-specific properties. Biochem J 389, 629–636 (2005).

11. K. G. Bache, A. Brech, A. Mehlum, H. Stenmark, Hrs regulates multivesicular body formation via ESCRT recruitment to endosomes. J Cell Biol 162, 435–442 (2003).

12. M. Komada, N. Kitamura, Growth factor-induced tyrosine phosphorylation of Hrs, a novel 115-kd protein with a structurally conserved putative zinc finger domain. Mol Cell Biol 15, 6213–6221 (1995).

13. K. G. Bache, C. Raiborg, A. Mehlum, I. H. Madshus, H. Stenmark, Phosphorylation of Hrs downstream of the epidermal growth factor receptor. Eur J Biochem 269, 3881–3887 (2002).

14. G. Prag et al., The Vps27/Hse1 complex is a GAT domain-based scaffold for ubiquitin-dependent sorting. Dev Cell 12, 973–986 (2007).

15. J. R. Mayers et al., ESCRT-0 assembles as a heterotetrameric complex on membranes and binds multiple ubiquitinylated cargoes simultaneously. J Biol Chem 286, 9636–9645 (2011).

16. Y. Amano et al., Hrs recognizes a hydrophobic amino acid cluster in cytokine receptors during ubiquitin-independent endosomal sorting. J Biol Chem 2011, 1 (2011).

17. Z. Marijanovic, J. Ragimbeau, K. G. Kumar, S. Y. Fuchs, S. Pellegrini, TYK2 activity promotes ligand-induced IFNAR1 proteolysis. Biochem J. 397, 31–38. (2006).

18. K. G. Kumar et al., Site-specific ubiquitination exposes a linear motif to promote interferon-α receptor endocytosis. J Cell Biol. 179, 935–950. (2007).

19. A. V. Villarino, Y. Kanno, J. J. O'Shea, Mechanisms and consequences of Jak-STAT signaling in the immune system. Nat Immunol 18, 374–384 (2017).

20. L. C. Platanias, Mechanisms of type-I-and type-II-interferon-mediated signalling. Nature reviews. Immunology 5, 375–386 (2005).

21. C. Thomas et al., Structural linkage between ligand discrimination and receptor activation by type I interferons. Cell 146, 621–632 (2011).

22. S. Lochte, S. Waichman, O. Beutel, C. You, J. Piehler, Live cell micropatterning reveals the dynamics of signaling complexes at the plasma membrane. J Cell Biol 207, 407–418 (2014).

23. L. B. Ivashkiv, L. T. Donlin, Regulation of type I interferon responses. Nature reviews. Immunology 14, 36–49 (2014).

24. H. Asao et al., Hrs is associated with STAM, a signal-transducing adaptor molecule. Its suppressive effect on cytokine-induced cell growth. J Biol Chem. 272, 32785–32791. (1997).

25. K. G. Bache, C. Raiborg, A. Mehlum, H. Stenmark, STAM and Hrs are subunits of a multivalent ubiquitin-binding complex on early endosomes. J Biol Chem. 278, 12513–12521. Epub 12003 Jan 12527. (2003).

26. O. Lohi, V. P. Lehto, STAM/EAST/Hbp adapter proteins-integrators of signalling pathways. FEBS Lett 508, 287–290 (2001).

27. T. Takeshita et al., STAM, signal transducing adaptor molecule, is associated with Janus kinases and involved in signaling for cell growth and c-myc induction. Immunity. 6, 449–457. (1997).

28. K. Endo et al., STAM2, a new member of the STAM family, binding to the Janus kinases. FEBS Lett. 477, 55–61. (2000).

29. E. Mizuno, K. Kawahata, A. Okamoto, N. Kitamura, M. Komada, Association with Hrs is required for the early endosomal localization, stability, and function of STAM. Journal of biochemistry 135, 385–396 (2004).

30. C. Raiborg, H. Stenmark, The ESCRT machinery in endosomal sorting of ubiquitylated membrane proteins. Nature 458, 445–452 (2009).

31. L. Christ, C. Raiborg, E. M. Wenzel, C. Campsteijn, H. Stenmark, Cellular Functions and Molecular Mechanisms of the ESCRT Membrane-Scission Machinery. Trends in biochemical sciences 42, 42–56 (2017).

32. O. Soderberg et al., Direct observation of individual endogenous protein complexes in situ by proximity ligation. Nat Methods 3, 995–1000 (2006).

33. C. Raiborg et al., FYVE and coiled-coil domains determine the specific localisation of Hrs to early endosomes. J Cell Sci 114, 2255–2263 (2001).

34. S. Christoforidis et al., Phosphatidylinositol-3-OH kinases are Rab5 effectors. Nat Cell Biol 1, 249–252 (1999).

35. I. J. Lodhi et al., Insulin stimulates phosphatidylinositol 3-phosphate production via the activation of Rab5. Mol Biol Cell 19, 2718–2728 (2008).

36. S. R. Elkin et al., Ikarugamycin: A Natural Product Inhibitor of Clathrin-Mediated Endocytosis. Traffic 17, 1139–1149 (2016).

37. C. S. Wegner et al., Ultrastructural characterization of giant endosomes induced by GTPase-deficient Rab5. Histochem Cell Biol 133, 41–55 (2010).

38. A. H. van Boxel-Dezaire, M. R. Rani, G. R. Stark, Complex modulation of cell type-specific signaling in response to type I interferons. Immunity 25, 361–372 (2006).

39. D. A. Jaitin et al., Inquiring into the differential action of interferons (IFNs): an IFN-alpha2 mutant with enhanced affinity to IFNAR1 is functionally similar to IFN-beta. Mol Cell Biol 26, 1888–1897 (2006).

40. L. M. Rodahl, S. Stuffers, V. H. Lobert, H. Stenmark, The role of ESCRT proteins in attenuation of cell signalling. Biochem Soc Trans 37, 137–142 (2009).

41. D. Chmiest et al., Spatiotemporal control of interferon-induced JAK/STAT signalling and gene transcription by the retromer complex. Nature communications 7, 13476 (2016).

42. C. M. Blouin et al., Glycosylation-Dependent IFN-gammaR Partitioning in Lipid and Actin Nanodomains Is Critical for JAK Activation. Cell 166, 920–934 (2016).

43. U. Muller et al., Functional role of type I and type II interferons in antiviral defense. Science 264, 1918–1921. (1994).

44. E. Kalie, D. A. Jaitin, R. Abramovich, G. Schreiber, An interferon alpha2 mutant optimized by phage display for IFNAR1 binding confers specifically enhanced antitumor activities. J Biol Chem 282, 11602–11611 (2007).

45. R. Bago et al., Characterization of VPS34-IN1, a selective inhibitor of Vps34, reveals that the phosphatidylinositol 3-phosphate-binding SGK3 protein kinase is a downstream target of class III phosphoinositide 3-kinase. Biochem J 463, 413–427 (2014).

