## Supplementary figures for "STAM Interaction with Hrs Controls JAK/STAT Activation by Interferon-α at the Early Endosome"

### Materials and Methods

Cell culture, transfection and drug treatment

Cells were grown at 37°C under 5% $CO_2$ in Dulbecco's modified Eagle's medium (DMEM) complemented with 10% FBS (v/v) (Gibco, Life Technologies). HeLa cells were grown in DMEM high glucose Glutamax (Gibco, Life Technologies) supplemented with 5 mM pyruvate (v/v) (Gibco, Life Technologies) and 1% penicillin-streptomycin (v/v) (Gibco, Life Technologies). Human retinal pigment epithelial cells (hTERT-RPE1) that express high levels of endogenous IFNAR1 and IFNAR2 subunits (*41*), were grown in DMEM/F12 Glutamax (Gibco, Life Technologies). Murine embryo fibroblasts from IFNAR1-knockout mice (MEF IFNAR1$^{-/-}$, (*43*) kind gift of Silvio Hemmi) were grown in DMEM high glucose Glutamax (Gibco, Life Technologies).

Plasmids were either electroporated or transfected using commercial kits of Lipofectamine® LTX with Plus™ Reagent (Invitrogen, Life Technologies), Lipofectamine® 3000 (Invitrogen, Life Technologies) or X-tremeGENE HP DNA Transfection Reagent (Roche) following manufacturers' instructions. Electroporation of cells was performed with a pulse at 220V and 975 µF with a Gene Pulser® II module (Bio-Rad). SiRNAs were transfected with HiPerFect kit (Qiagen) following the manufacturer's instructions, and cells were incubated for three days before further experimentations. Depletion efficiency was assessed by immunoblotting.

Except when stated otherwise, human IFN-$\alpha$2 and IFN-$\beta$1a (kind gift of Pierre Eid) were used at 1000 U/ml. The IFN-$\alpha$ YNS mutant (kind gift of Gideon Schreiber) was described in (*44*). Alexa Fluor® 546 transferrin (Thermo Fisher) was used at 10 µg/ml. VPS34-IN1 is a selective inhibitor of the class III PI3K Vps34 (*45*) and was purchased from the University of Dundee (D. Alessi laboratory). HeLa cells were treated with 30 µM during 40 min at 37°C. Ikarugamycin (Sigma) was diluted in DMSO and used at 4 µM after 20 min of preincubation.

Plasmids, and antibodies

DNA constructs were verified by sequencing. For expression in eukaryotic cells, human IFN-$\alpha$R1 WT and IFN-$\alpha$R1 K501/525/526R (Ub1/2) cDNAs were cloned in pcDNA3.1-Flag-zeo. Human Hrs was cloned in a pIRES-puro2C-mCherry empty vector to obtain Hrs-mCherry tagged protein. Human TYK2 was cloned in a pmCherry-C vector to obtain TYK2-mCherry tagged fusion protein. STAM2A/Hbp was subcloned from pMIW-HA-Hbp to pEGFP-N2 (Clontech), pEGFP-C1 (Clontech) or pmCherry-N1 (Clontech) to obtain respectively eGFP-STAM2A, STAM2A-eGFP or mCherry-STAM2A. In this study, we used murine Hbp for investigating human STAM2A as they both share 86% of amino acids identity. pEF-IRES-IFN-$\alpha$R2, and pcAGGS-IFNAR1-GFP WT were kind gifts respectively of P. Eid and M. Fukata, peGFP-rab5Q79L of S. Miserey-Lenkei, peGFP-FYVE of P. Gleeson and RFP-rab5/CAAX Q79L of A. Saltiel.

Primary antibodies were mouse anti-phospho-STAT1 Tyr 701 (BD Transduction Laboratories, # 612132, 1:2000 for western blot and 1:200 for immunofluorescence), rabbit anti-STAT1 (Cell Signaling, # 9172, 1:1000 for western blot), rabbit anti-phospho-JAK1 Tyr 1022/1023 (Cell Signaling, # 3331, 1:1000 for western blot), rabbit anti-TYK2 (UpState, number 06-638, 1:1000 for western blot), rabbit anti-phospho-TYK2 Tyr 1054/1055 (Cell Signaling Technology, # 9321, 1:1000 for western blot), mouse anti-Phosphotyrosine PY20 (BD Transduction Laboratories, # 610000, 1:1000 for western blot), rabbit anti-IFNAR1 (Abcam, # ab124764, 1:2000 for western blot and 1:200 for immunofluorescence), mouse anti-IFNAR1 AA3 (Biogene, kind gift of Darren Baker, 1:100 for immunofluorescence and FACS, 1:1000 for immunoprecipitation), mouse anti-IFNAR1 64G12 (kind gift of Pierre Eid, 1:1000 for western blot), rabbit anti-Phospho-IFNAR1 Tyr 466 (Santa Cruz Biotechnology, # sc-13114-R, 1:1000 for western blot), mouse anti-Alpha-tubulin (Sigma, clone B512, T5168, 1:5000 for western blot), mouse anti-clathrin heavy chain (BD Transduction Laboratories, # 610500, 1:5000 for western blot), goat anti-EEA1 (Santa Cruz Biotechnology, clone N19, #sc-6415, 1:100 for immunofluorescence), rabbit anti-Hrs (Abcam, #ab155539, 1:200 for immunofluorescence and 1:1000 for immunoprecipitation), rabbit anti-Hrs 958/3 (from Sylvie Urbé, 1:1000 for western blot), rabbit anti-phospho-Hrs Tyr 334 842/3 (from Sylvie Urbé, 1:1000 for western blot), rabbit anti-STAM1/2 (Abcam, # ab76061, 1:100 for immunofluorescence, 1:2500 for western blot), mouse anti-biotin (BioLegend, # 409002, 1:150 for FACS), Alexa Fluor® 647 streptavidin antibody (ThermoFisher Scientific, # S32357, 1 :100 for FACS), rabbit anti-eGFP and rabbit anti-mCherry (recombinant antibody platform, Institut Curie, Paris, France, 1:1000 for western blot). Secondary antibodies conjugated to Alexa 488, Cy3, Cy5, AMCA or HRP (Beckman Coulter, Jackson ImmunoResearch and Invitrogen, 1:200 for immunofluorescence, 1:5000 for western blot).

RNA interference

Except when stated otherwise, siRNA were used at 20 nM and shRNA at 20 μg. Depletion efficiency was assessed by immunoblotting. Control siRNA (Dharmacon, Thermofisher, # SI03650325, AATTCTCCGAACGTGTCACGT), Hrs pool of 4 siRNA FlexiTube GeneSolution (Dharmacon, Thermofisher, L-016835-00, GAGGUAAACGUCCGUAACA,GCACGUCUUUCCAGAAUUC,AAAGAACUGUGGC CAGACA,GAACCCACACGUCGCCUUG), siSTAM1 (Eurogenetec, GGCUUUGACUCUUCUAGGA) used at 20 nM, siSTAM2 (Eurogenetec, GGAGCGAAAGAUUGCCUAA) used at 20 nM for targeting both STAM2A and STAM2B genes, siJAK1 (Qiagen, # SI00605514, ACCGGATGAGGTTCTATTTCA), siTYK2 (Qiagen, # SI0222321, CCAUCUGGUAAUAAACUCATT), CHC pool of 4 siRNA (Qiagen, # GS1213) each used at 5 nM. pSuper-scramble and pSuper-shHrs (5'GATCCCCCGACAAGAACCCACACGTCTTCAAGAGAGACGTGTGGGTTCTTGT CGTTTTTGGAAA-3'; kind gift of Philippe Benaroch).

Cell stimulation by IFNs

Cells were treated with or without 1000 U/ml IFN-α or IFN-β at 37°C for the indicated times. For biochemical analysis, cells were washed with PBS and lysed in sample buffer (62.5 mM Tris/HCl pH 6.0, 2% v/v SDS, 10% glycerol, 40mM dithiothreitol and 0.03% w/v phenol red). Total lysates were analyzed by SDS-PAGE and Western blot analysis and immunoblotted with the indicated antibodies. Chemiluminescence detection was performed with SuperSignal West Dura Extended Duration Substrate or with SuperSignal West Femto Substrate (Thermo Scientific Life Technologies). Signal acquisition and quantification were done on a ChemiDoc MP Imaging System (Bio-rad). Phosphorylated and total forms of the proteins were quantified and normalized to clathrin heavy chain or tubulin levels in the same lysate. Phosphorylated protein over total ratio was determined for each time point.

Phospho-STAT1 nuclear translocation

For immunofluorescent analysis, cells were grown on coverslips, treated as described above, and then fixed with cold methanol at -20°C for 10 min. After washing with 0.2% BSA (w/v) in PBS, cells were incubated with primary anti-pSTAT1 antibody for 1 h at room temperature and revealed by using a Cy3-conjugated goat anti-mouse secondary antibody. Coverslips were mounted in Fluoromount-G mounting medium (eBioscience) with 2 μg/ml DAPI (Sigma-Aldrich) to counterstain nuclei. Pictures were acquired on a Leica DM 6000B inverted widefield microscope equipped with a HCX PL Apo 63X NA 1.40 oil immersion objective and an EMCCD camera (Photometrics CoolSMAP HQ). Nuclear translocation was quantified with a homemade plugin on ImageJ software (NIH) by calculating the nucleo-cytoplasmic ratio of phospho-STAT1 signal (nuclei masks were realized with DAPI staining).

IFNAR1 endocytosis assay

Cells grown on coverslips were incubated on ice with anti-IFNAR1 antibody for 30 min, washed in ice-cold PBS and then incubated in DMEM/F12 at 37°C for the indicated times. Cells were fixed in 4% PFA in PBS for 10 min at room temperature, quenched in 50 mM $NH_4Cl$ for 10 min and permeabilized with 0.05% saponin in 0.2% BSA in PBS for 20 min. Immunostainings were performed with the indicated antibodies. Pictures were acquired with a Nikon A1R confocal microscope equipped with a CFI Plan Apo VC 60X NA 1.4 oil immersion objective.

Fluorescence-activated cell sorting (FACS)

RPE1 cells were grown on a 6-well plate for 72 h and transfected with the indicated siRNA. To monitor endocytosis of endogenous IFNAR1, cells were incubated on ice with the anti-AA3 antibody for 30 min, washed in ice-cold PBS and then incubated in DMEM/F12 at 37°C with or without IFN-α or IFN-β for 10 min. To stop endocytosis, cells were washed in cold PBS, and then resuspended by Accutase for 2 min at 37°C. Cells were washed by 3% FBS (w/v) in PBS at 4°C and centrifuged at 800 g for 3 min. The supernatant was discarded and these two steps were repeated twice. Cells were fixed with 4% PFA at 4°C for 10 min, quenched in 50 mM $NH_4Cl$ for 10 min. Cells were not permeabilized to stain only the remaining pool of cell surface IFNAR1 after internalization. After washing with 3% FBS (w/v) in PBS for 15 min at 4°C, cells were first incubated in suspension with a primary anti-biotin antibody for 45 min on ice and then, with a streptavidin Alexa647-conjugated secondary antibody in order to amplify the AA3 signal. The intensity of fluorescence of extracellular IFNAR1 was measured by the BD AccuriTM C6 flow cytometer (BD Biosciences) and data analyzed with BD AccuriTM software.

Proximity Ligation Assay (PLA)

PLA kit was purchased from Sigma-Aldrich, and the assay was performed according to the manufacturer's protocol. PLA experiments were performed in combination with IFNAR1 endocytosis assay or directly after treatment with IFNs. In the former condition, RPE1 cells grown on coverslips were incubated on ice with anti-IFNAR1 antibody for 30 min, washed in ice-cold PBS and then incubated with or without IFNs in DMEM/12 at 37°C for the indicated time. Cells were fixed in 4% PFA in PBS for 10 min at room temperature, quenched in 50 mM $NH_4Cl$ for 10 min and permeabilized with 0.1% Triton X100 (w/v) for 10 min. Cells were then incubated with the primary antibody against Hrs or STAM as indicated. After 1 hour of incubation, cells were washed in PBS containing 0.2% (w/v) BSA and secondary antibodies conjugated with PLA probe MINUS and PLUS were added to the reaction for 1 h at 37°C, which allows ligation to occur if the two oligonucleotides are less than 40 nm to each other. The signal was then amplified using fluorescently labeled oligonucleotides in association with a polymerase that allows to visualize distinct fluorescent dots.

Co-immunoprecipitation

Cells were lysed in 1% NP-40 in TNE (10 mM Tris/HCl pH 7.5, 150 mM NaCl, 0.5 mM EDTA) with protease inhibitors cocktail (Roche) for 30 min at 4°C. Cleared lysates (16,000g, 10 min, 4°C) were incubated overnight at 4°C under rotation with 1 µg/ml of the indicated antibody followed by incubation for 1 h with 25 µl of protein A/G magnetic beads (Thermo Scientific) in the case of endogenous proteins. For tagged proteins, 25 µl of GFP-Trap or RFP-Trap beads (Chromotek) were used. After three washes in TNE, immunoprecipitated beads were eluted following the manufacturers' instructions.

FLIM FRET

FLIM measurements were done in frequency domain by phase modulation on a custom system based on a commercial module (Lifa, Lambert Instruments) coupled to a Nikon TE 2000 TIRF inverted microscope equipped with a Coolsnap HQ CCD camera (Photometrics). A 473 nm modulated laser diode (Omicron) was used for excitation of the donor fluorophore and a 100x 1.49 NA TIRF objective. The laser light was coupled using a Yokogawa CSU10 spinning-disk module. Fluorescence emission was selected by a band-pass filter (500-550 nm). Transfected RPE1 cells were imaged by widefield illumination microscopy by a mercury lamp with standard Nikon filter cubes and a Coolsnap Ez CCD camera (Photometrics) prior to FLIM measurements in order to estimate the expression level of both GFP- and mCherry-tagged proteins. The measurements of eGFP-STAM fluorescence lifetime were restricted to early endosomes, as determined by widefield illumination microscopy for EEA1 positive staining, in cells expressing or not the fluorescent fusion protein Tyk2-mCherry. All the experiments were performed at 37°C with 5% $CO_2$. FLIM images were analyzed on area containing GFP-positive structures with the LI-FLIM software (Lambert Instruments) in order to determine the mean fluorescence lifetime for each structure. Time exposure was comprised between 200-1000 msec.

Statistical analysis

Statistics were calculated using Prism 6.0 software (GraphPad Software).

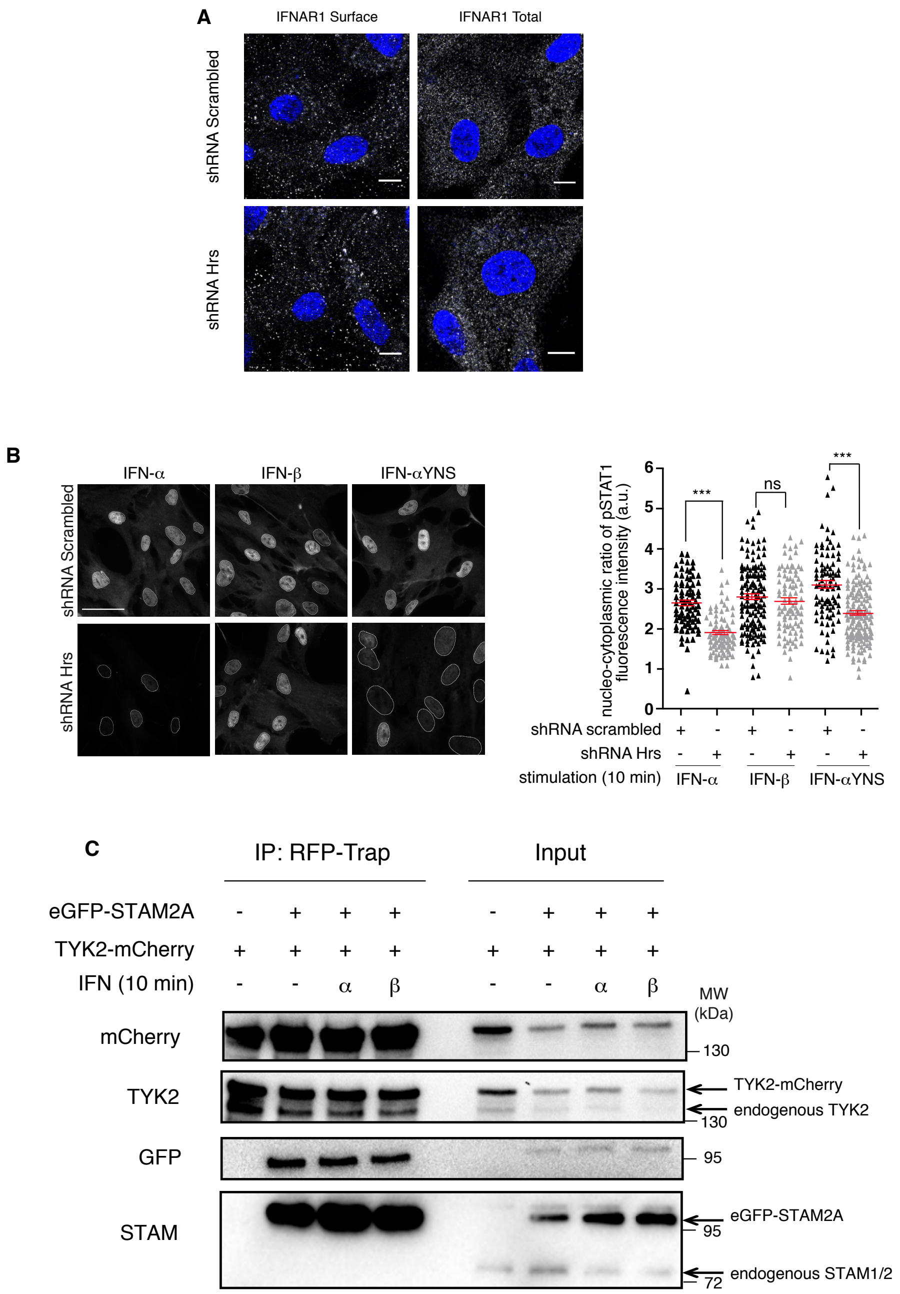


**Fig. S1. Additional experiments on interaction studies and IFN selectivity.**

(**A**) Expression levels of surface (left panels) and total (right panels) of IFNAR1 subunits in control (upper panels) or Hrs depleted RPE1 cells (lower panels) analyzed by confocal microscopy showing no effect of Hrs depletion on IFNAR1 expression. Scale bar 10 μm. (**B**) Left, nuclear translocation of pSTAT1 in cells depleted or not from Hrs upon stimulation with IFN-α, IFN-β, and IFN-α YNS for 10 min. Right, the nucleus/cytosol fluorescence ratio for pSTAT1 is quantified from the fluorescence imaging. P-values were calculated with one-way ANOVA with Kruskal-Wallis multiple comparison test. ****P*<0.001; ns, not significant. Results are representative of 2 independent experiments. Scale bar, 50 μm. (**C**) RFP-Trap immunoprecipitation of TYK2-mCherry in RPE1 cells transfected with TYK2-mCherry and eGFP-STAM2A after stimulation or not with IFN-α or IFN-β for 10 min as indicated. Data are representative of 2 independent experiments.

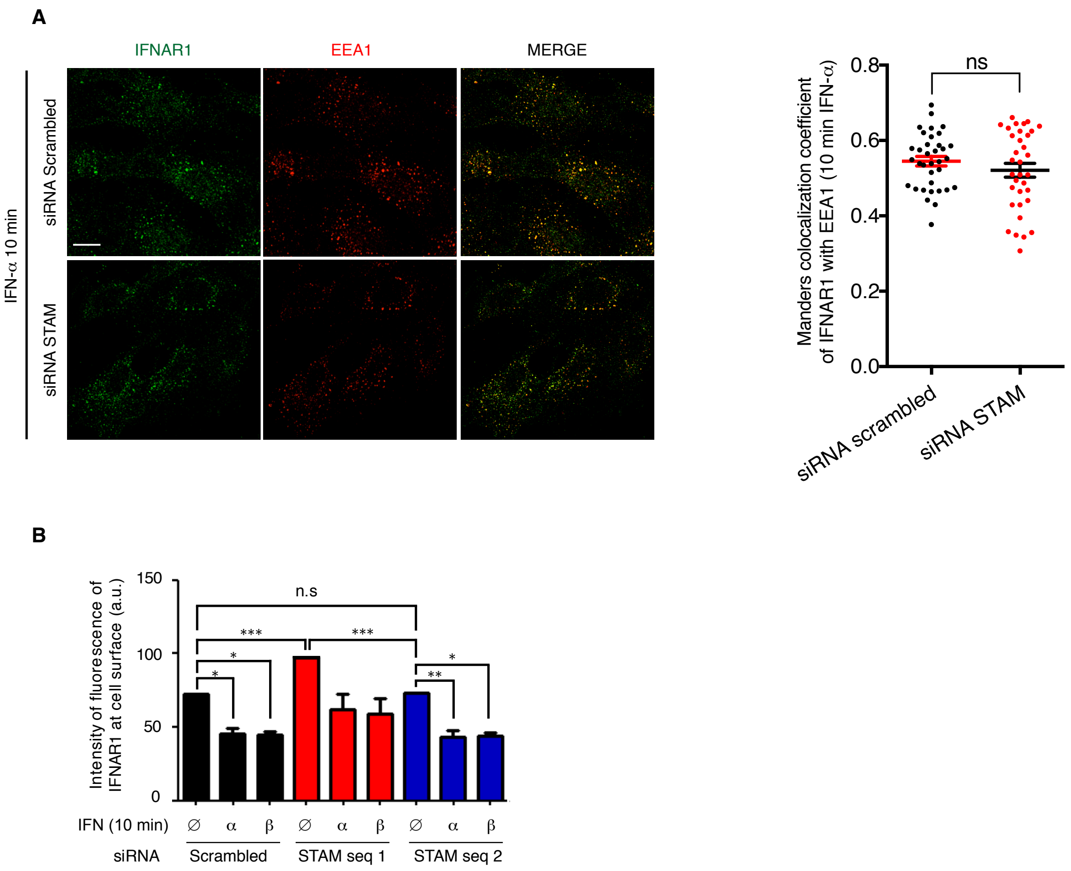


**Fig. S2. STAM is not required for IFNAR1 endocytosis.**

(**A**) IFNAR1 uptake in control (siRNA scrambled) or STAM depleted RPE1 cells showing that IFNAR1 trafficking is not impaired upon STAM depletion. Cells were stimulated for 10 min with IFN-α. Following fixation, cells were co-labeled for EEA1 and analyzed by confocal microscopy. Scale bar 10 μm. Right, quantification of co-localizations are expressed as the Manders' coefficient, indicating the proportion of IFNAR1 pixels containing EEA1 pixels in 3 independent experiments. (**B**) Quantification of IFNAR1 surface staining monitored by FACS in control or STAM depleted RPE1 cells stimulated with IFN-α or IFN-β for 10 min. Data are representative of 2 independent experiments.

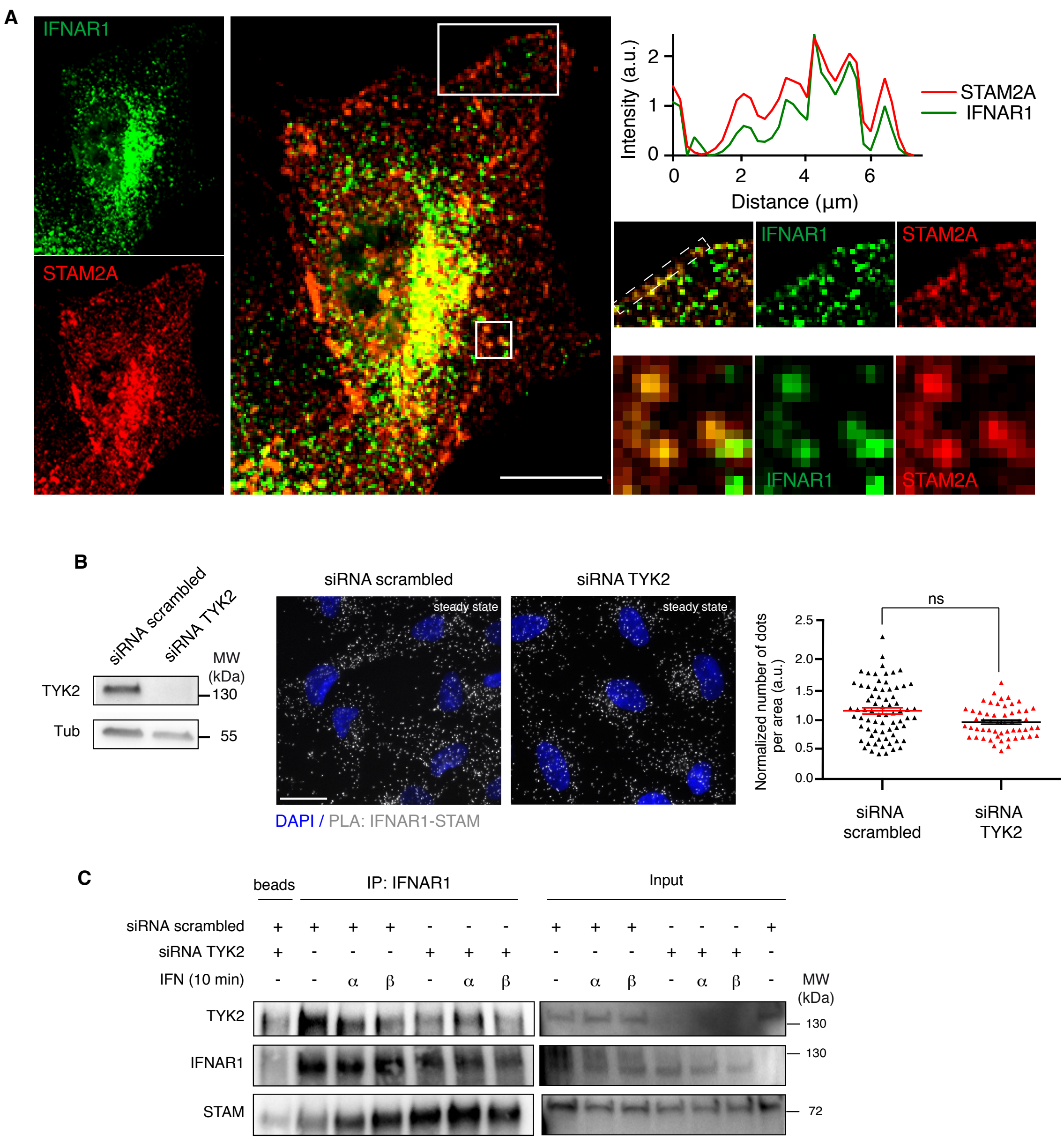


**Fig. S3. Additional experiments on IFNAR1 and STAM interaction.**

(**A**) Co-localization of endogenous IFNAR1 and STAM2A by fluorescent confocal microscopy in RPE1 cells at steady state. Magnification (upper right) and profile of IFNAR1 and STAM colocalization plot at the plasma membrane from the dashed line insert. Scale bar 10 µm. (**B**) Left, immunoblots for TYK2 expression levels in RPE1 cells transfected or not with siRNA TYK2 for 72 h. Middle, PLA experiments monitoring the interaction between total IFNAR1 and STAM in control or TYK2 depleted RPE1 cells. Right, corresponding quantifications show mean values ± SEM. Statistical significance was determined using a two-tailed t test. ns: no significant. Scale bar 10 µm. (**C**) Co-immunoprecipitation of endogenous STAM with endogenous IFNAR1 subunit in control or TYK2 depleted RPE1 cells treated or not for 10 min with IFN-α or IFN-β as indicated. Data are representative of 3 independent experiments.

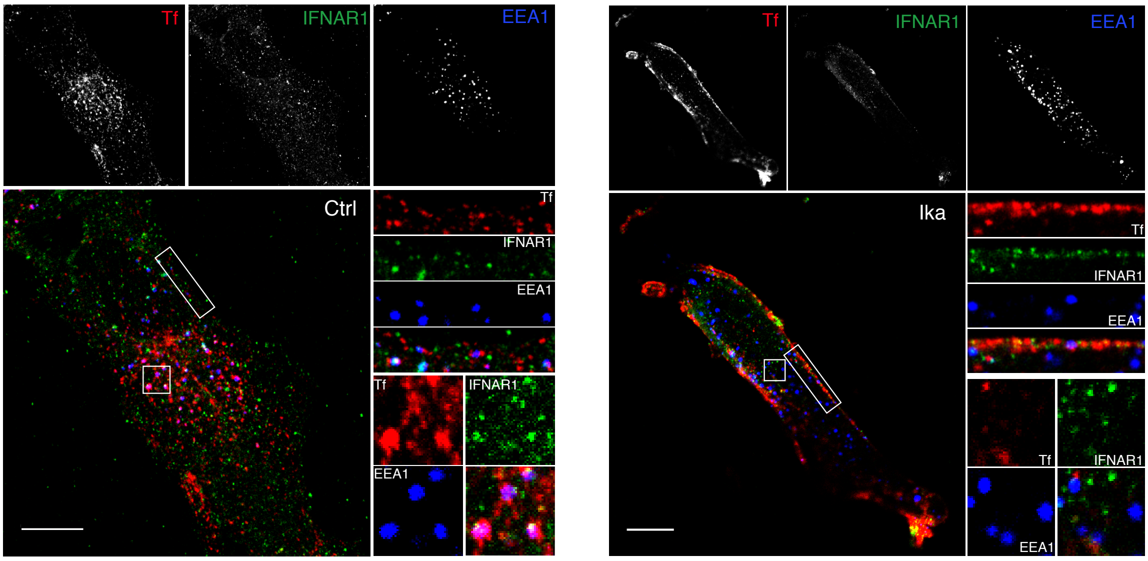


**Fig. S4. Ikarugamycin inhibits clathrin-dependent endocytosis of IFNAR and Transferrin.** RPE1 cells treated with 4 μM ikarugamycin (Ika) or not (Ctrl) were incubated for 20 min at 4°C with Alexa Fluor 546 transferrin, anti-IFNAR1 antibody and stimulated with IFN-α. Cells were then shifted to 37°C for 10 min to allow endocytosis, fixed, immunostained for EEA1 and observed by confocal microscopy. Data are representative of 3 independent experiments. Scale bars, 10 μm.

**Author contributions:** N.Z., C.M.B. and C.V. designed the study, performed the experiments and analysis of the results. N.Z., C.M.B., C.V. and C.L. wrote the manuscript. D.C. and L.J. assisted with experiments and provided several reagents. C.L. supervised and directed the research. All authors discussed the results and commented on the manuscript.
